# microRNA tissue atlas of the malaria mosquito *Anopheles gambiae*

**DOI:** 10.1101/177964

**Authors:** Lena Lampe, Elena A. Levashina

## Abstract

*Anopheles gambiae* mosquitoes transmit the human malaria parasite *Plasmodium falciparum*, which causes the majority of fatal malaria cases worldwide. The hematophagous life style defines the mosquito reproductive biology and is exploited by *P. falciparum* for its own sexual reproduction and transmission. The two main phases of the mosquito reproductive cycle, pre-vitellogenic (PV) and post-blood meal (PBM) shape its capacity to transmit malaria. Transition between these phases is tightly coordinated to ensure homeostasis between mosquito tissues and successful reproduction. One layer of control is provided by microRNAs, well known regulators of blood meal digestion and egg development in *Aedes* mosquitoes. Here, we report a global overview of tissue-specific miRNA expression during the PV and PBM phases and identify miRNAs regulated during PV to PBM transition. The observed coordinated changes in the expression levels of a set of miRNAs in the energy-storing tissues suggest a role in the regulation of blood meal-induced metabolic changes.

## INTRODUCTION

*Anopheles gambiae* females are major vectors of the human malaria parasite *Plasmodium falciparum* in sub-Saharan Africa. Mosquito females require a blood meal for egg production, which the parasite exploits as a route of transmission and development, thus linking the mosquito infection and its reproductive (also called gonotrophic) cycle. As reproduction is central to every organism, the hematophagic life style directs mosquito behavior, physiology, and metabolism. The gonotrophic cycle is divided into two phases – the pre-vitellogenic (PV) and post blood meal (PBM) phase. During the PV phase, egg maturation (vitellogenesis) is arrested at a premature stage until the blood meal-mediated uptake of amino acids and lipids (1–3). The PV phase, maintained by the juvenile hormone (JH), prepares the carbohydrate-consuming female mosquitoes for blood feeding by accumulating energy resources in the fat body, the major storage tissue in insects, and by activating their host-seeking behavior (4–9). Successful blood feeding initiates the PBM phase, which is driven by the steroid hormone ecdysone (10,11). Amino acids, released into the mosquito circulatory system after blood meal digestion, activate the target of rapamycin (TOR) pathway (12–14). TOR together with ecdysone, synthesized from the blood-borne cholesterol, trigger expression of lipid transporters such as lipophorin and vitellogenin that deliver lipids to the developing ovaries (2,12,15,16). Within 48 h, a blood meal is fully digested, nutrients absorbed, and oogenesis is completed. However, a blood meal may also bring *Plasmodium* infective stages or gametocytes that fuse in the mosquito midgut to produce motile ookinetes (17). The ookinetes traverse the midgut epithelium at 18-24 h post BM and establish an infection at the basal side of the midgut wall (17). Here within the next ten days, the parasites undergo replication and maturation of the infective forms (18). Disruption of the gonotrophic cycle by ecdysone analogs or by restriction of lipid trafficking decreases *Plasmodium* development (19–22). Therefore, the PBM phase shapes both mosquito reproduction and *P. falciparum* development, two crucial components of malaria transmission. Repeated blood feedings increase female reproductive fitness by boosting its nutritional resources and promote *P. falciparum* development by providing key nutrients and a route for transmission to the next host.

The metabolic transition from the PV to PBM phase entails a complex regulatory network of hormonal, transcriptional and post-transcriptional changes in multiple tissues. However, the mechanisms of tissue homeostasis during this transition remain incompletely understood. Recently, microRNAs (miRNAs) emerged as powerful regulators of developmental and metabolic switches. miRNAs fine tune JH and ecdysone pulses and, thereby, shape developmental transitions in the fruit flies (23–25). In mosquito adults, miR-8, miR-309, miR-275, miR-1174 and miR-1890 have been implicated in the regulation of PBM (26–31), and miR-305 in parasite development (32). However in *A. gambiae*, functional studies were performed only for three out of 180 miRNAs identified by bioinformatics and experimental approaches (31,33–35). The functional diversity of miRNAs is further amplified by the capacity of each miRNA locus to generate two miRNAs arms, which differ in their seed sequence and target distinct sets of mRNAs. Several studies observed a bias in arm selection for different species and tissues, suggesting that it might represent an additional layer of miRNA regulation (33,36–38). In addition, the same miRNA can regulate distinct processes in a tissue-specific manner by binding different targets. For example, the *Drosophila* miR-8 regulates the production of myogenic peptide hormone in the fat body, whereas it controls synapse structure in the brain (39,40). So far, all studies on *A. gambiae* miRNAs focused either on whole mosquitoes or on few selected tissues (26,31–35). However, a comprehensive overview of tissue-specific miRNA expression would be beneficial for further characterization of miRNA function during PV to PBM transition.

Here, we report a global overview of miRNA expression in the head, midgut, abdominal carcass (the fat body) and ovaries of *A. gambiae* females during the transition from the PV to PBM phase, and after *P. falciparum*-infection. We show that the majority of mosquito miRNAs are tissue-specific, except for a small cluster of ubiquitously expressed miRNAs. Further, we identify four miRNAs, whose expression levels are regulated by blood feeding in the fat body, ovaries, and midgut. Our study provides the first comprehensive miRNA tissue atlas of the female *A. gambiae* mosquito during the gonotrophic cycle and identifies miRNAs with potential roles in this critical process.

## METHODS

### Mosquitoes

We used *Anopheles coluzzii* Ngousso (*TEP1*S1*) mosquitoes. Mosquito lines differ in the genotype of *TEP1*, the major mosquito immune marker. Ngousso line was initially isolated as a mixed population of *TEP1*S1*, *\*S2* and *\*S1/S2* genotypes. To reduce background genetic variation, we selected to work with *TEP1*S1* homozygous line to avoid potential genetic interferences. Mosquitoes were reared at 30°C 80% humidity at 12/12 h day/night cycle with a half-hour long dawn/dusk periods. All mosquitoes were fed *ad libitum* with 10% sugar solution.

### Blood meal and *P. falciparum* infections

Blood meals and *P. falciparum* infections were performed using membrane feeders (41,42). To establish gametocyte cultures, *Pf* NF54 asexual cultures (parasitaemia >2%) were harvested by centrifugation for 5 min at 1,500 rpm, washed with fresh red blood cells and diluted to 1% total parasitaemia in complete gametocyte medium at 4% hematocrit. Gametocyte cultures were incubated at 37°C with 3% O^2^ and 4% CO^2^. Medium was changed daily for 15-16 days on heated plates to reduce temperature drops. On day 14 after establishment, (i) gametocytaemia was checked by Giemsa stain of gametocyte cultures-smears, and (ii) parasite exflagellation rates were estimated by microscopy.

Mosquitoes were fed with either uninfected or *P. falciparum*-infected blood using an artificial feeder system as described in Lensen *et al.* (41,42). The feeding system was prepared by covering the bottom of the midi-feeders with a stretched parafilm. The midi-feeder was then attached to a 37°C water bath system, allowing water flow through the feeder. The blood (Haema, Berlin, Germany) was introduced to the feeder and mosquitoes were fed for 15 min. Unfed mosquitoes were removed and only fully engorged females were kept for further analyses.

### Sample collection

#### Microarray and qPCR

Three-day-old virgin females were divided into three groups and fed with: (i) 10% sugar (sugar-fed; PV phase); (ii) human blood (blood-fed; PBM phase); (iii) *P. falciparum*-infected blood (infected; PBM phase). After feeding, all groups were kept at 26°C 80% humidity. At 18 h post feeding, 15 females per group were dissected on ice and the heads, midguts, ovaries and abdominal carcasses were pooled according to the tissue. The samples were homogenized in TRIzol using a beat-beater (Qiagen) at 50 rpm. Homogenized samples were kept at −80°C until further usage.

#### Time course

For miRNA expression kinetics, 15 unfed and 15 blood-fed females were dissected on ice at 20, 24, 28, 32 and 48 h after blood meal. The heads, midguts, ovaries and abdominal carcasses were pooled according to the tissue, immediately homogenized in TRIzol and kept at −80°C for RNA isolation.

### RNA isolation

#### Microarray

Total RNA was isolated using the miRNeasy Mini Kit (Qiagen) according to the manufacturer’s recommendations. Briefly, after phase separation by centrifugation total RNA was purified by silica membrane, eluted with water and stored at −80°C. The total RNA yield was measured with a NanoDropND-1000 Spectrophotometer. The integrity of total RNA was assessed with a 2100 Bioanalyzer and a RNA 6000 Nano LabChip kit (Agilent). Furthermore, the ratio of miRNA/small RNAs from isolated total RNA was monitored by the Agilent Small RNA kit.

#### qPCR

Total RNA was isolated by TRIzol according to the manufacturer’s recommendations. The total RNA yield was measured with a NanoDropND-1000 spectrophotometer.

### Microarray analyses

Custom 8-plex 60K mosquito miRNA microarrays (Design Name: Agilent-049943, ID Name: Custom_Mosquito_miRNA, Design Format: IS-62976-8-V2, AMADID 016436) (Agilent) were used for one-color hybridizations. The microarray included miRNAs identified in *Apis mellifera*, *Anopheles gambiae*, *Tribolium castaneum*, *Culex quinquefasciatus*, *Aedes aegypti*, *Bombyx mori* and *Drosophila melanogaster.* Total RNA (100 ng) was processed with the miRNA Complete Labeling and Hyb Kit (Agilent) according the supplier’s recommendations. In brief, samples were dephosphorylated with Calf Intestine Alkaline Phosphatase and labeled with Cy3 in a T4 RNA ligase-mediated reaction with 3’, 5’-cytidine bisphosphate (Cy3-pCp). The labeling reaction was column-purified, vacuum-dried using a speed-vac at 45°C and resuspended in a blocking and hybridization buffer. After hybridization, the microarrays were washed, scanned at 5 µm resolution with a G2565CA high-resolution laser microarray scanner, and features were extracted. Results were analyzed by the R limma package. Briefly, signal intensities were corrected for the background using the normexp function and quantile normalized. All spot replicates were pooled using the avereps function producing an Elist containing log2 relative expression data. Values of the negative control spots were subtracted from the mean intensities and all negative probes were removed. Expression values were calculated as mean values of the dipteran miRNA expression of the four biological replicates. As the microarray contained miRNAs of multiple insect species including miRNA orthologues, only probes with at least 14 nucleotides complementarity to the annotated *A. gambiae* miRNAs were included in further analysis. In total, 506 microarray probes were included, representing 86 mature *A. gambiae* miRNAs annotated in the miRBase (Table S1). Differentially regulated miRNAs were identified by fitting a linear regression model to the data and empirical Bayes statistics on the model in R. All microarray probes that differ in sequence were analyzed and visualized separately. Finally, we aimed to provide an overview about miRNA enrichment in mosquito tissues. To this end, we set an enrichment cut-off of 1.5-fold (Fig. 6). Two studies in *A. gambiae* and *Ae. aegypti* showed that miR-1175-3p is a midgut-specific miRNA (26,35). We took expression levels of this miRNA across different tissues and observed a 1.5-fold higher expression of miR-1175 in the midgut compared to other tissues. Therefore, we applied this cut-off to other miRNAs.

### cDNA synthesis and qPCR

Total RNA was reverse-transcribed with the miScript Kit (Qiagen). Expression levels of mature miRNAs and of mRNAs were measured using the Quantitect SYBR Green PCR Kit (Qiagen). Relative quantities of miRNA expression were normalized to the gene encoding ribosomal protein S7 (*RPS7*). The miRNA primers were obtained as miRNA Primer Assay (Qiagen).

## RESULTS

We set out to develop a global overview of tissue-specific miRNA expression in the head, midgut, ovary and fat body, during the gonotrophic cycle of *A. gambiae* and after *P. falciparum* infections. To this end, we determined miRNA expression levels using the Agilent custom-designed miRNA microarray in the three groups of females fed with: (1) sugar; (2) human blood or (3) *P. falciparum*-infected blood (Fig. 1).

**Figure 1:**
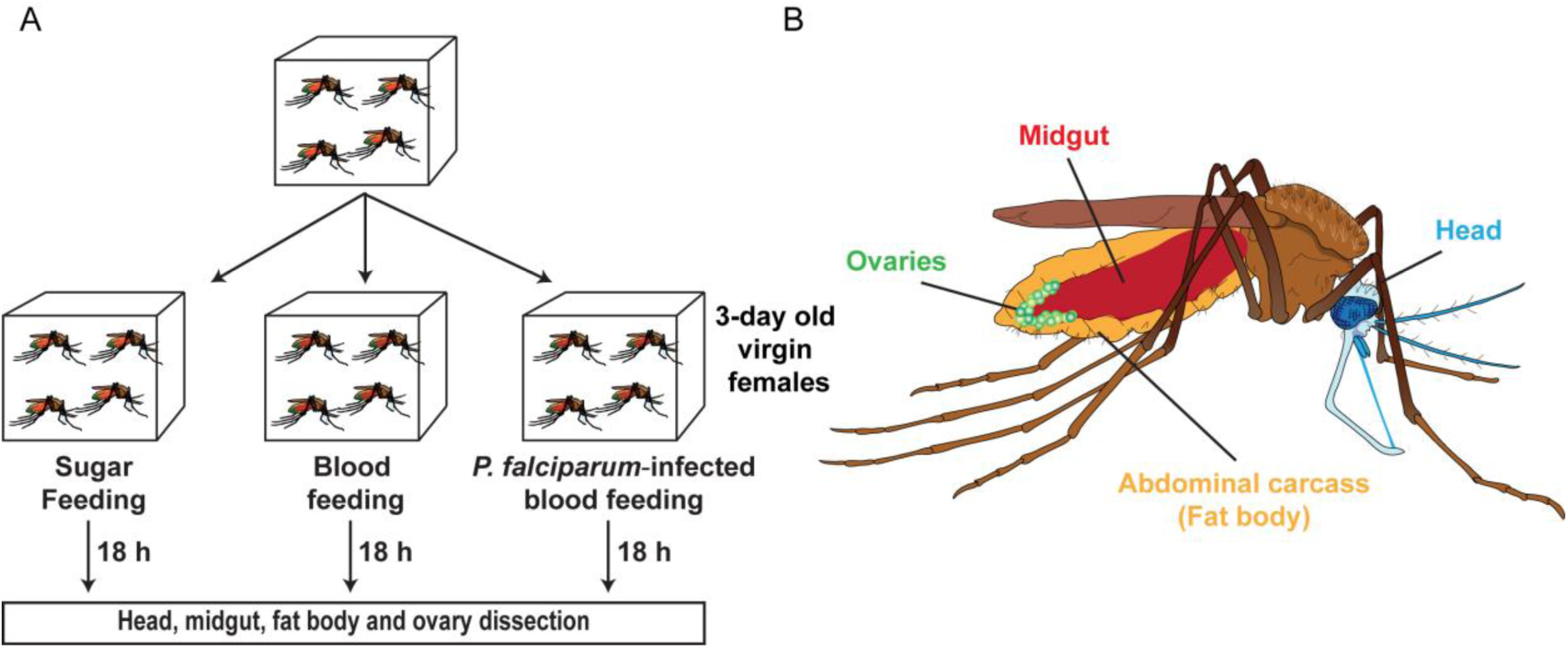
Experimental set up for tissue-specific miRNA expression analysis. (A) Mosquitoes were fed with: (1) 10% sugar, (2) blood or (3) *P. falciparum*-infected blood. At 18 h post blood feeding, mosquitoes from each group were dissected. (B) The head, midgut, fat body and ovaries were examined for tissue-specific miRNA expression.

### MicroRNA expression in the pre-vitellogenic phase

We first examined expression levels of miRNAs in the pre-vitellogenic (PV) phase. Microarray analysis detected expression of 74 out of 86 miRNA in at least one tissue. The microarray probes matched almost all *A. gambiae* miRNAs annotated in the miRBase, as well as the highly conserved insect miRNAs (Table S1). One limitation of the microarray approach is that it is applicable only to known miRNAs. Instead, RNA-seq technologies are better suited for discovery of novel miRNAs and for overcoming problems of cross-kingdom miRNA hybridization in blood-fed mosquitoes. The problem of RNA-seq is its quantitative inaccuracy as the majority of the newly-identified *Anopheles*-specific miRNAs reported by previous studies are expressed at very low levels (6,7). Therefore, our microarray approach covers the more highly expressed as well as conserved miRNAs. High levels of expression across all tissues were detected only for miR-8, whereas nine miRNAs (miR-1, miR-2, let-7, miR-34, miR-125, miR-184, miR-277, miR-306, and miR-957) had moderate to high levels of expression in at least two tissues (Fig. 2). A second cluster of 18 miRNAs included tissue-specific miRNAs with intermediate expression levels. The largest number of tissue-specific miRNAs (cut-off 1.5-fold) was observed in the head (17 miRNAs), including the well-known brain-enriched miR-7 and miR-124. Three miRNAs, miR-281, miR-1174 and miR-1175, were enriched in the midgut, whereas miR-10, miR-92b, miR-279, miR-989, and miR-998 were predominantly detected in the ovaries. Surprisingly, not a single miRNA was exclusively expressed in the fat body. Microarray results were further validated by independent quantitative PCR (Fig. 3A-D).

**Figure 2:**
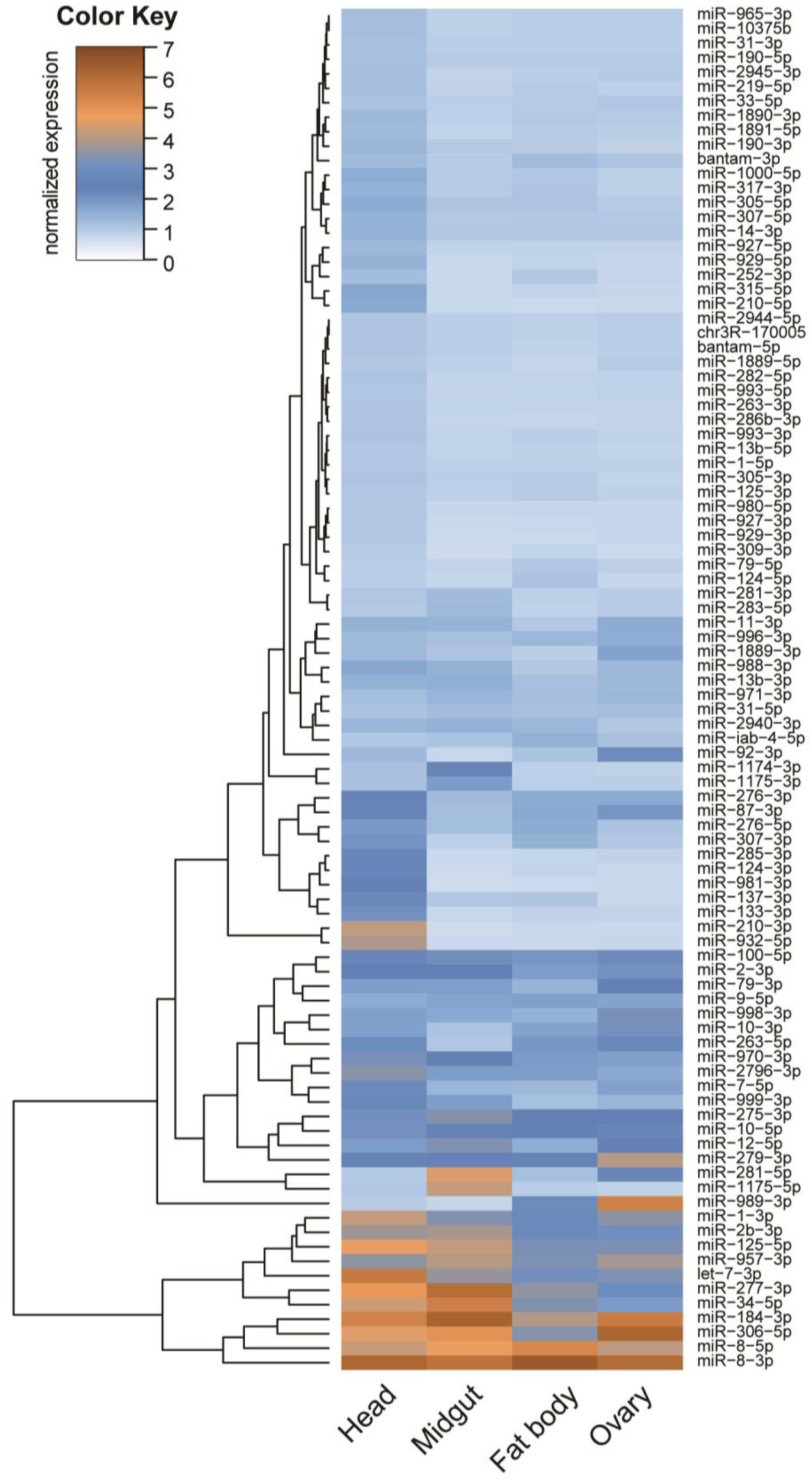
Tissue-specific miRNA expression in the 3-day-old *A. gambiae* female. Heatmap of miRNAs expression levels in the head, midgut, fat body and ovaries of sugar-fed mosquitoes in the PV phase. Colour gradient from light blue to dark brown represents an increase in miRNA expression.

**Figure 3:**
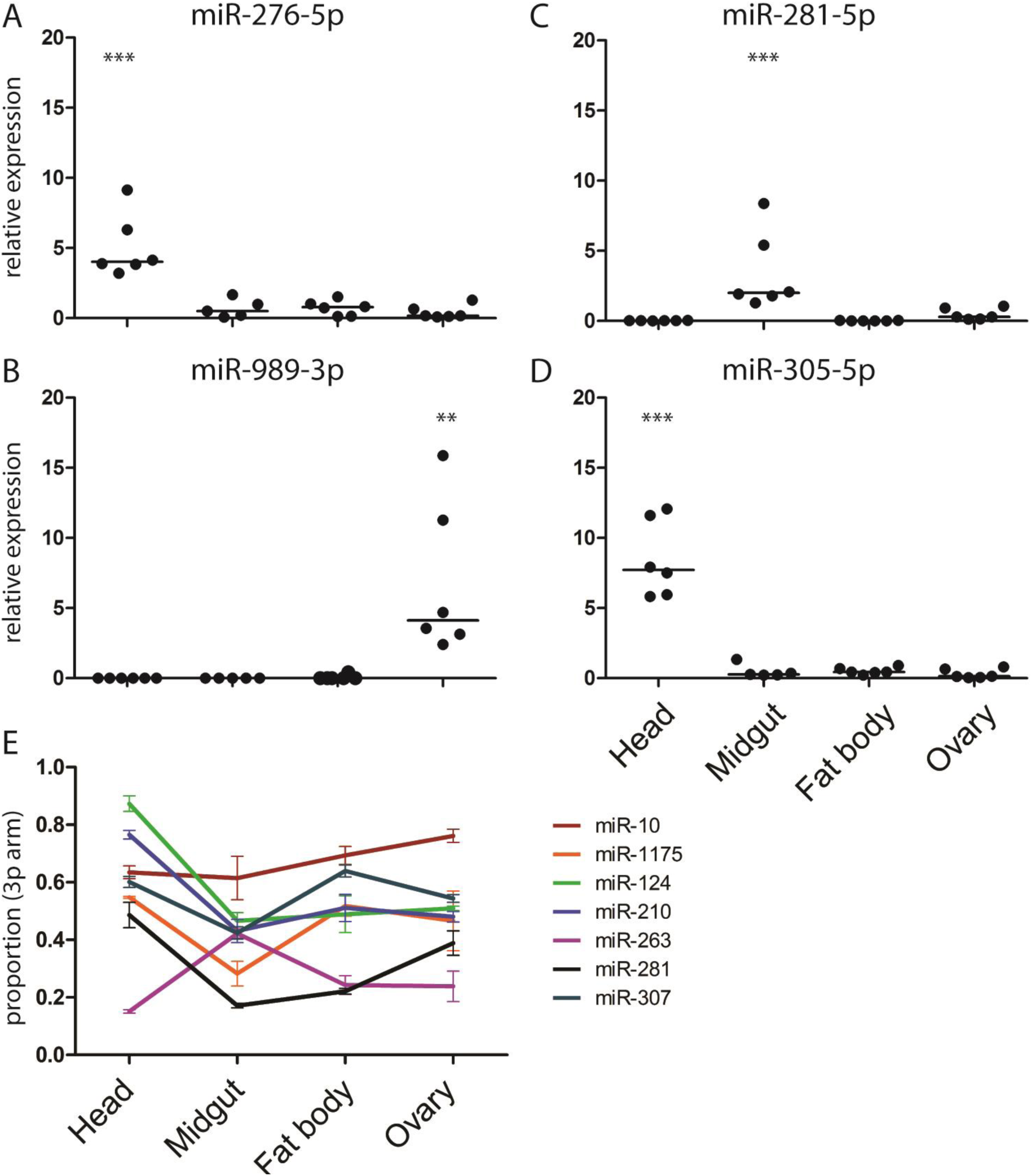
Tissue*-specific expression of miRNAs quantified by qPCR* of (A) miR-276-5p, (B) miR-281-5p, (C) miR-989-3p and (D) miR-305-5p. miRNA expression values are plotted as dots with the line indicating the mean. miRNA expression levels were normalized using the ribosomal protein *RPS7* gene. Results of six independent experiments are shown. *\*\*\* p<0.0005 and ** p<0.005,* one-way ANOVA followed by Tukey’s post-hoc test. (E) Tissue-specific differences in miRNA 3p-arm proportion. All depicted 3p-arm proportions significantly change between tissues tested by one-way ANOVA (p-value < 0.05; Table S3).

The mature miRNA can be generated from the 3p or 5p arm of the pre-miRNA. We used arm-specific probes to quantify tissue-specific expression of miRNA arms and identified 20 miRNAs expressing both arms (Table S2). Among those, seven miRNAs showed tissue-dependent arm bias (Fig. 3E). Head-specific enrichment in the 3p arm was observed for miR-124 and miR-210, whereas the 5p arm bias was detected for miR-1175 in the midgut and for miR-263 in the head, fat body and ovaries. Furthermore, higher levels of the 5p arm were detected in the head and midgut for miR-10. A complex pattern of arm usage was observed for miR-281 and miR-307. While the miR-281-5p arm was enriched in the fat body and midgut, expression of the 3p arm increased in the head and ovary. For miR-307, the 3p arm was predominantly expressed in the head and fat body. We concluded that during the PV phase, expression of the majority of the miRNAs is tissue-specific and, in some cases, is regulated at the arm level.

### Tissue-specific miRNA expression after blood feeding and after *P. falciparum* infection

We next examined miRNA expression during the PBM phase and after *P. falciparum* infection at 18 h post feeding (hpb) (Fig. S1). At this time point, all midgut samples from blood-fed females had high background levels, probably due to unspecific binding of human short RNAs and DNAs present in the blood meal. Therefore, we excluded these samples from further analyses. Interestingly, blood feeding impacted expression levels of very few miRNAs in other tissues. Major changes were observed in the fat body, where blood feeding increased levels of miR-275-3p, miR-276-5p and miR-305-5p, and reduced levels of miR-989-3p (Fig. 4A). Higher levels of miR-305-5p were also detected in the ovaries (Fig. 4B). No changes in miRNA levels after blood meal were observed in the head (Fig. 4C). Surprisingly, infections with *P. falciparum* had no effect on miRNA expression in any tested tissue (Fig. S2A-C). The expressional changes detected by microarray analysis were further confirmed by qPCR, except for miR-989-3p with very low expression levels in the fat body (Fig. 4D). Taken together, our results identified three miRNAs whose expression was modulated during PV to PBM transition in the fat body, whereas expression of miR-305-5p was also upregulated in the ovaries.

**Figure 4:**
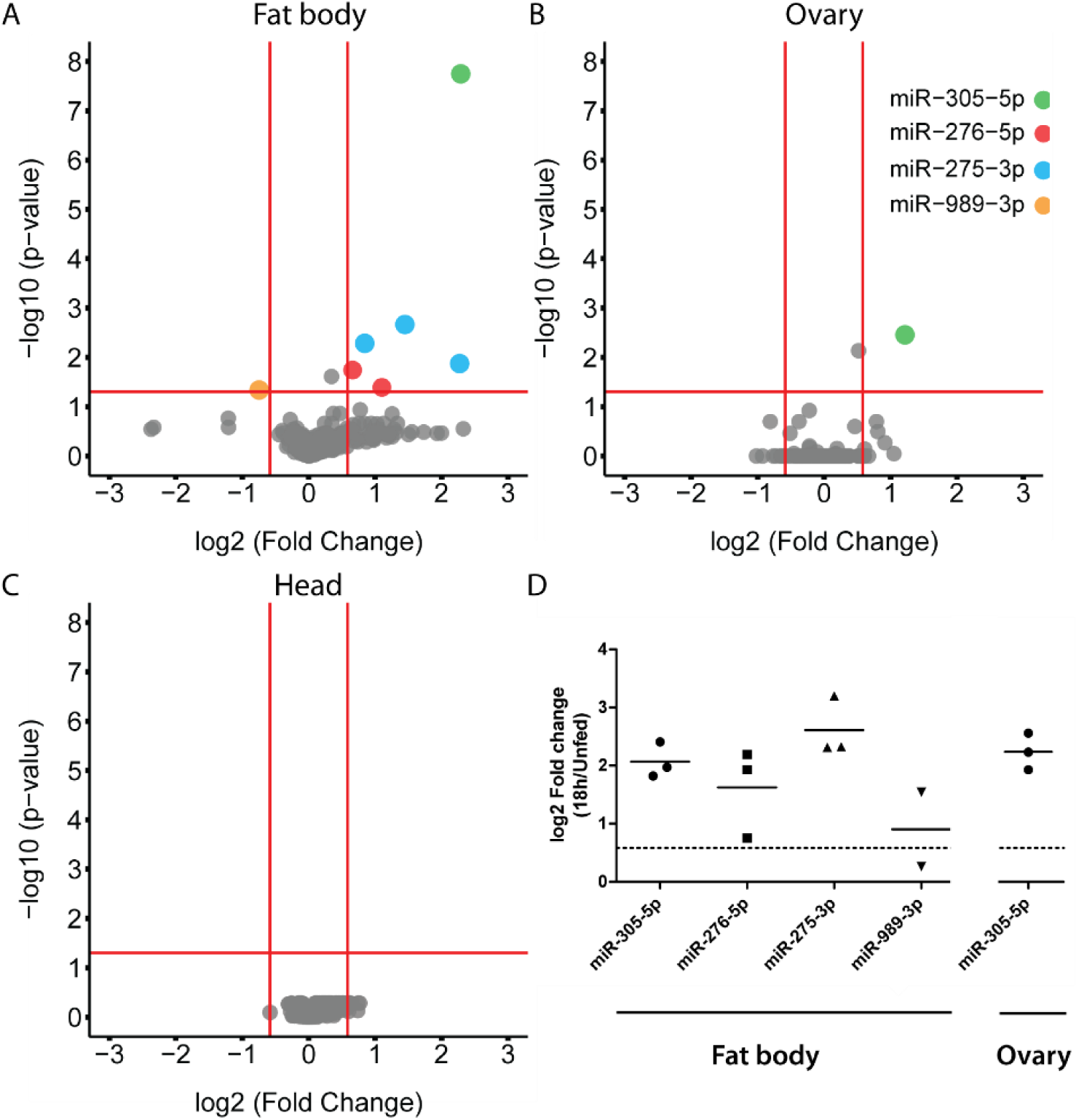
Differentially expressed miRNAs at 18 h post blood feeding (hpb) in: (A) the fat body, (B) the ovaries and (C) the head. Cut-off log10 p-value of +/−1.3 (p<0.05) and log2 fold change of +/−0.58 (FC>1.5) (n=4) were used to detect significant changes in miRNA expression by fitting a linear model with empirical bayes statistics. The dots represent different miRNA probes. (D) Blood feeding-induced changes (log2 FC) in the fat body and ovary of miRNA expression levels at 18 hpb measured by qPCR. Results of three independent experiments are shown, horizontal lines indicate mean values.

### Expression of blood meal-induced miRNAs during the PBM phase

The kinetics of tissue-specific expression of the blood feeding-induced miRNAs was further analyzed by qPCR. The highest expression levels of miR-305-5p, miR-275-3p and miR-276-5p were observed in the head before blood feeding, while expression of miR-989-3p was only detected in the ovaries (Fig. 5A). Expression levels of miR-275-3p, miR-276-5p and miR-305-5p transiently declined in the head at 24 and 28 hpb and regained the initial levels by 32 hpb (Fig. 5A). In contrast, high levels of these miRNAs were detected in the fat body and midgut at the same time points (Fig. 5B and 5C). In the fat body, miR-275-3p and miR-276-5p levels peaked at 28 and 32 hpb and remained high at 48 hpb. High inter-replicate variation was observed for miR-305-5p whose levels increased between 20-48 hpb (Fig. 5B). In the midgut, miR-275-3p, miR-276-5p and miR-305-5p peaked at 24 to 32 hpb, but in contrast to the fat body, their levels slightly declined by 48h post BM (Fig. 5C). High expression levels of miR-989-3p detected in the ovaries before blood feeding, gradually declined during 48 hpb. In contrast, low transcript levels of miR-305-5p, miR-275-3p and miRNA-276-5p in the ovaries slightly increased at 28 and 32 hpb (Fig. 5D). We concluded that PV to PBM transition regulates miRNA expression levels in a tissue- and time-specific manner.

**Figure 5:**
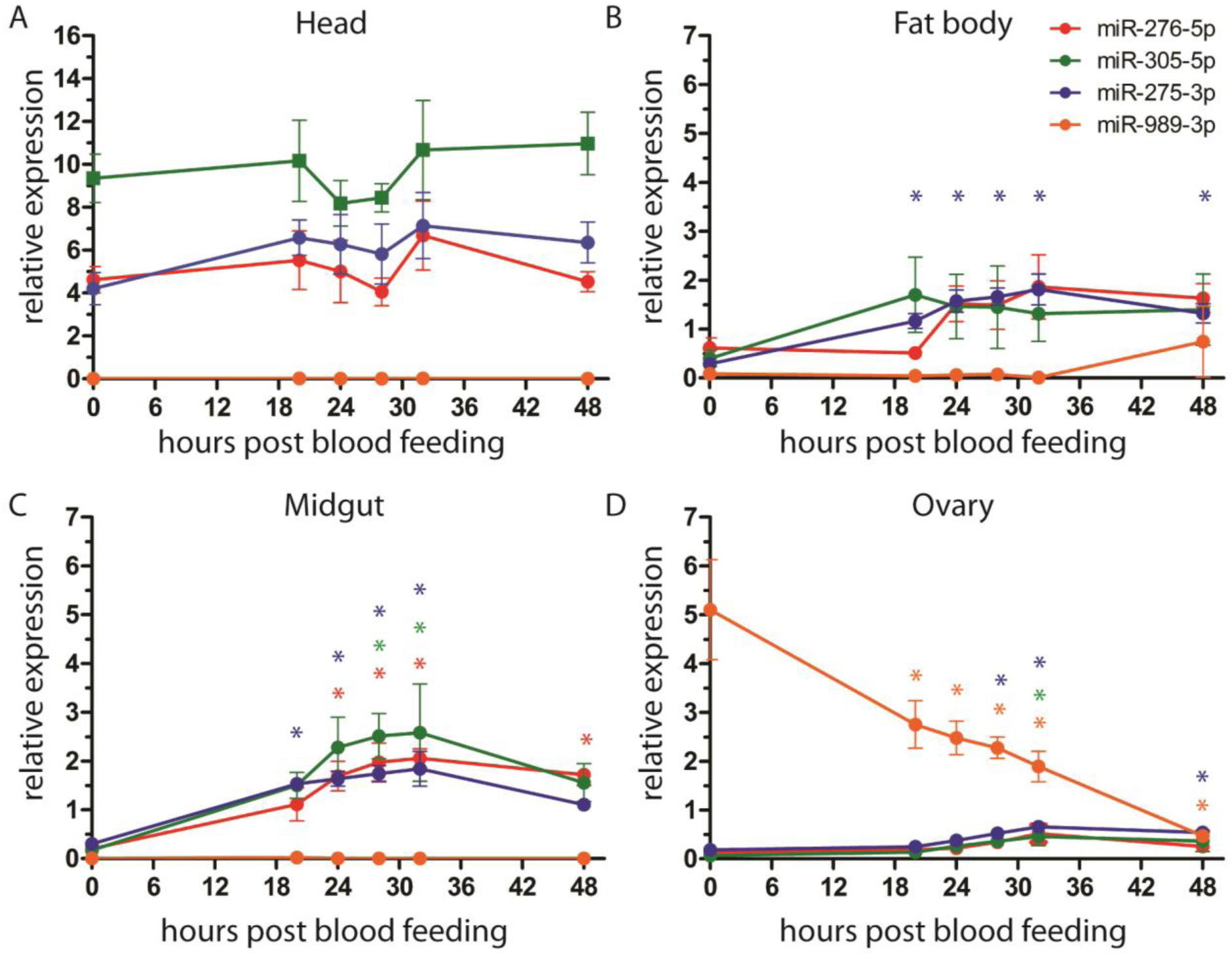
Tissue-specific expression of miR-276-5p, miR-305-5p, miR-275-3p and miR-989-3p during the PBM phase in (A) the head, (B) the fat body, (C) the midgut and (D) the ovary. Results of three independent experiments are shown as mean +/−SEM. For statistical analyses all time points were compared to 0 h (unfed) by one-way ANOVA, Tukey’s post-hoc test and significant differences are shown by asterisk (* = p<0.05; n=5).

## DISCUSSION

We report a global overview of miRNA expression in the *A. gambiae* tissues during the late PV and PBM phases. We found that the majority of *A. gambiae* miRNAs are expressed in a tissue-specific manner with the exception of the ubiquitously highly expressed miR-8. While six miRNAs showed tissue-specific bias in arm expression, one third of the miRNAs expressed both arms. Previous RNA sequencing studies of whole mosquitoes identified miR-10, miR-184, miR-263, miR-281 and miR-306 as the most abundant miRNAs in adult females (33,34). We show that expression of these abundant miRNAs is restricted to specific tissues (Fig. 6A). These results are in agreement with functional specialization of tissues, which is in part regulated by miRNAs (23,40,43).

**Figure 6:**
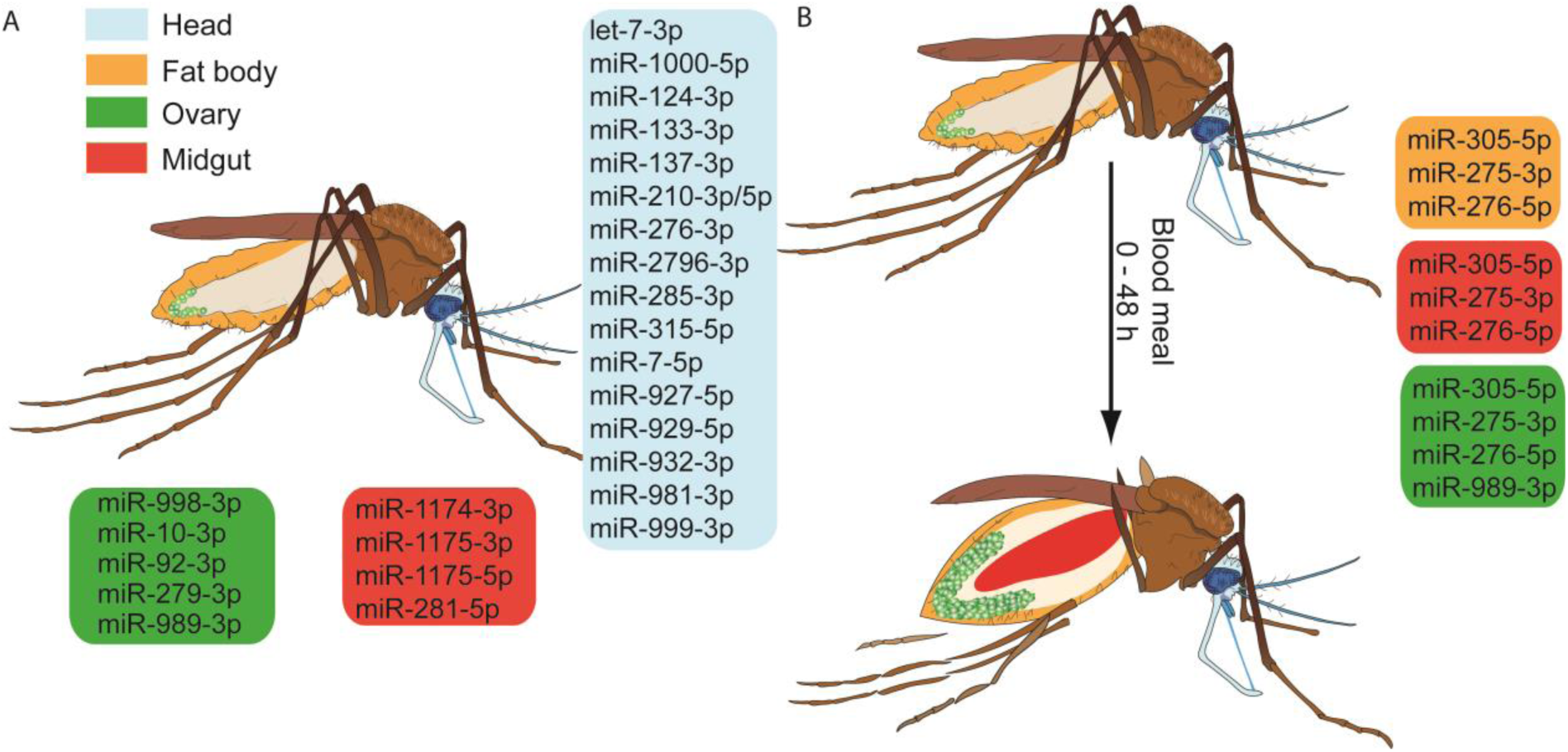
Schematic summary of tissue-specific miRNA expression during the gonotrophic cycle of the female *A. gambiae* mosquito. (A) Tissue-enriched miRNAs in sugar-fed mosquitoes. Enrichment cut-off for tissue-specific miRNAs was set at 1.5-fold. (B) miRNAs regulated during the PBM phase. The coloured boxes show tissue-specific miRNAs (blue = head, yellow = fat body, green = ovary, red = midgut).

Reproduction in many mosquito species is tightly coupled to blood feeding. During the PV phase, females feed on sugars to increase energy reserves for efficient egg development (4,5). Previous reports showed that miR-1174 regulates sugar absorption in the midgut of *Aedes* female mosquitoes (26). We demonstrate conservation of the midgut-specific expression of miR-1174/1175 and miR-281 between *A. gambiae* and *Ae. aegypti* (26,35,44). In addition, we also observed high levels of miR-277 in the midgut. *Drosophila* miR-277 regulates branched-chain amino acid (BCAA) catabolism and activates TOR signaling in the thoracic muscles (45).

BCAAs serve as signaling metabolites in tissue communication, systemically promote anabolic metabolism and release hormones from the intestinal tract in mice (46–49). It is possible, that BCAA catabolism in the midgut signals food uptake to distant tissues. Further investigations of miR-277 function in the midgut are needed to explore an interesting link between food digestion, TOR signaling and systemic metabolic changes. TOR signaling is tightly connected to synthesis of JH, the major driver of the PV phase (50). During insect development, systemic JH levels serve as a checkpoint for sufficient energy reserves and body size before transition to the next developmental stage. *Drosophila* miR-2 generates a threshold for developmental transitions by regulating downstream JH signaling (51). The rise of JH levels with nutrient availability during the PV phase suggests that the JH checkpoint function may be also conserved in mosquitoes (8,9,52). Furthermore, high expression levels of miR-2 in the PV phase indicate that expression of this miRNA may also regulate JH signaling in *A. gambiae*.

Brain tissues, especially the central nervous system, play essential roles in the PV phase by regulating mosquito olfactory and circadian behaviors (53,54). Efficient response to blood feeding depends on timely accumulation of the ovary ecdysteroidogenic hormone in the granules of neurosecretory cells (55). We found that the majority of the tissue-specific miRNAs are expressed in the head, including the well-known miRNAs that regulate olfactory sensing and circadian rhythms (56–60). In mosquitoes, these processes shape such important vector competence traits as host seeking and biting behavior, therefore, further investigation of the head-specific miRNAs should identify new factors and mechanisms that regulate vector competence.

Although feeding induces massive physiological changes in the mosquito, we identified only modest changes in expression levels of miRNAs during the early PBM phase in the head, fat body and ovaries (Fig. 6B). Note that no information could be generated for the midgut tissues pending the technical problem of cross-hybridization. Interestingly, blood meal-induced miRNAs in the fat body follow similar expression patterns in the midgut indicating potential roles of these miRNAs in the regulation of mosquito metabolism. In line with this hypothesis, miR-275 and miR-305 modulate metabolic processes in other insects. In *Ae. aegypti*, miR-275 ensures successful blood meal digestion, fluid excretion and, consequently, egg development (27). In agreement with miR-275 expression patterns in *A. gambiae*, TOR and ecdysone control expression of this miRNA in *Ae. aegypti* (27). Similarly, TOR and insulin pathways in *Drosophila* regulate expression of miR-305 in the fat body and the gut, respectively, where miR-305 fosters adaptation to starvation by balancing stem cell renewal and differentiation in a nutrient-dependent context (61,62). Interestingly, increased levels of miR-305 in the midgut 24 h post *P. falciparum* infection negatively impact parasite development in *A. gambiae* (32). Although the mechanisms of parasite inhibition are currently unknown, it is plausible that miR-305 affects *Plasmodium* development by regulating midgut homeostasis and/or nutrient availability. Further studies should examine whether the effect of miR-305 on *Plasmodium* development is midgut-specific. We did not detect any miRNAs regulated by *P. falciparum* infection in the head, fat body and ovaries during the early PBM phase. Whether parasite replication regulates miRNA expression during later stages of the PBM phase remains to be investigated.

As ovary growth and maturation is initiated in the second half of the PBM phase, it is not surprising that we detected only minor changes in the expression levels of ovarian miRNAs. Indeed, at 24 h post blood feeding the oocytes only begin to accumulate lipids, necessary for their maturation. Nevertheless, the observed down regulation of miR-989 coincides in time with initiation of border cell migration, which in *Drosophila* is regulated by miR-989 (63). Therefore, *A. gambaie* miR-989 may have a conserved role in induction of oogenesis.

Collectively, our results show tissue-specific expression patterns of 74 *A. gambiae* miRNAs during the PV and PBM phases. Recent metabolic studies during the gonotrophic cycle in *Aedes* mosquitoes identified highest metabolic activity at 36 hpb in the fat body (4,5). Therefore, we propose that tissue-specific modulation of miRNA expression by blood feeding reported in this study may contribute to regulation of metabolic changes during the mosquito gonotrophic cycle.

## Acknowledgments

The authors thank Liane Spohr and Sandrina Koppitz for mosquito breeding as well as Dana Tschierske and Daniel Eyermann for cultivation of the *P. falciparum* parasite. We thank Prof. Hilary Ranson, Dr. Craig Wilding and Dr. Inna Biryukova for sharing the custom microarray design and we acknowledge the technical support of Dr. Hans-Joachim Mollenkopf and Ina Wagner from the microarray core facility at the Max Planck Institute for Infection Biology. Finally, we are also grateful to Dr. Maiara Severo for comments on the manuscript.

---

**Figure S1:**
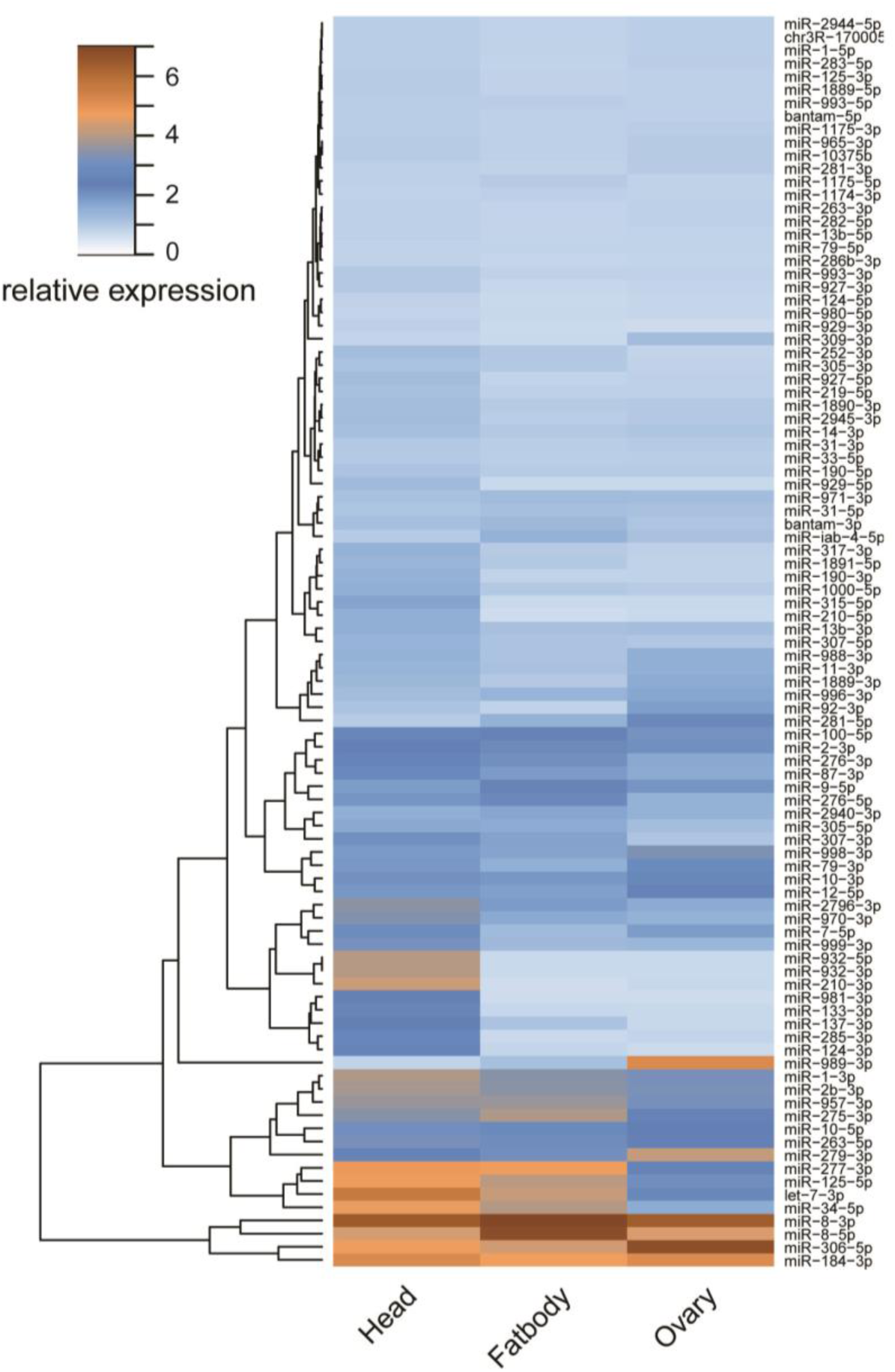
Tissue-specific miRNA expression in blood-fed *A. gambiae*. Heatmap of miRNA expression levels in the head, midgut, fat body and ovaries at 18 h post blood meal. Colour gradient from light blue to dark brown represents the increase in miRNA expression.

**Figure S2:**
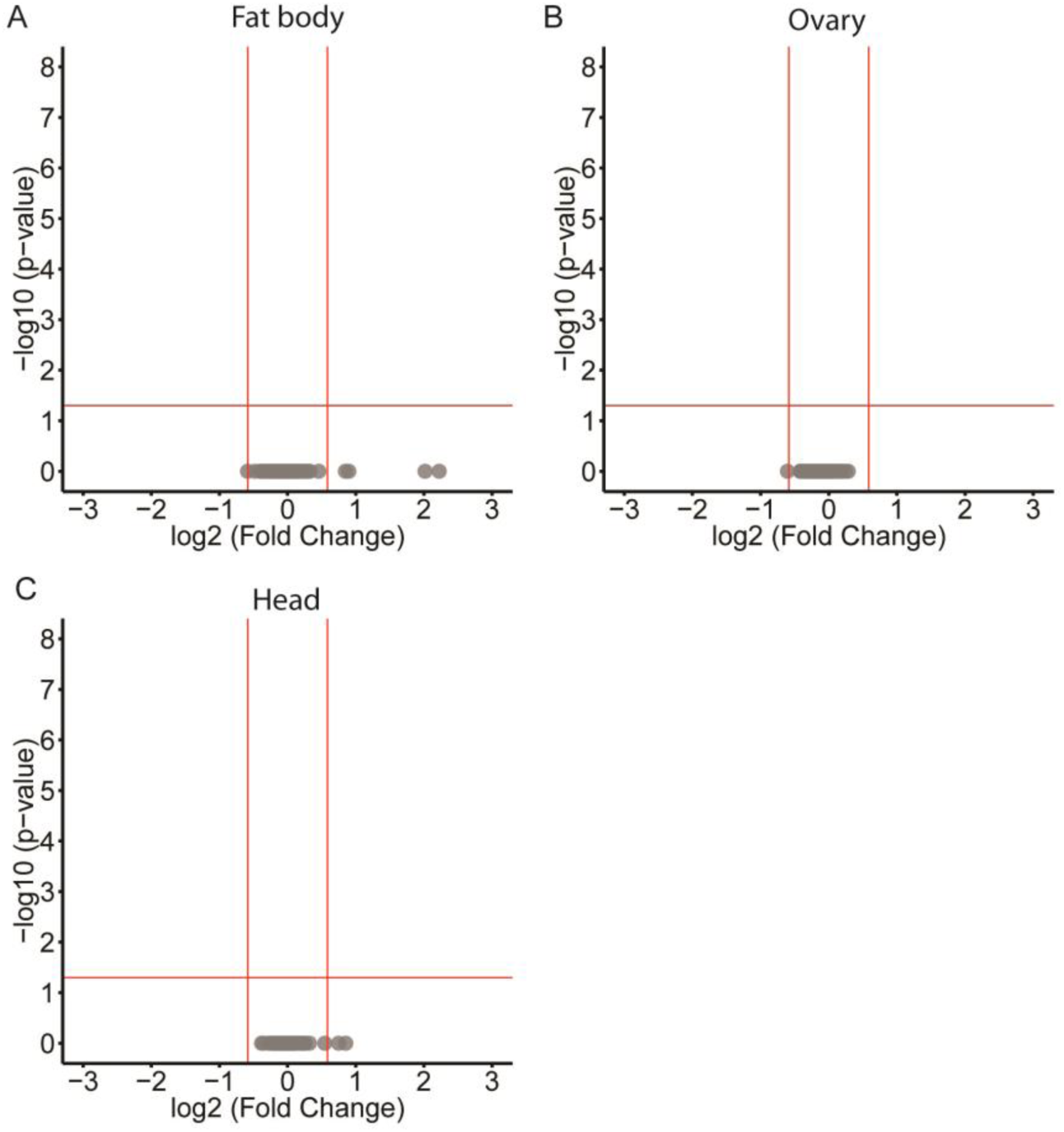
Differentially expressed miRNAs at 18 h post *P. falciparum*-infected blood feeding in *A. gambiae* tissues. (A) fat body, (B) ovaries and (C) head. Cut-off log10 p-value of +/−1.3 (p<0.05) and log2 fold change of +/−0.58 (fold-change>1.5) (n=4) were used to detect significant differences in miRNA expression.

**Table S1:** The microarray repertoire of *A. gambiae* miRNA arms

|  |  |  |
| --- | --- | --- |
| aga-miR-212 | aga-miR-79-5p | aga-miR-278-3p |
| aga-miR-929-5p | aga-miR-87-3p | aga-miR-9c-5p |
| aga-miR-100-5p | aga-mir-2796-5p | aga-miR-9a-5p |
| aga-miR-281-5p | aga-mir-2940-5p | aga-miR-957-3p |
| aga-miR-1175-5p | aga-mir-33-3p | aga-miR-1890-3p |
| aga-miR-308-3p | aga-miR-263b-3p | aga-miR-317-3p |
| aga-miR-282-5p | aga-miR-1174-5p | aga-mir-286b-5p |
| aga-miR-92b-3p | aga-miR-988-5p | aga-mir-2945-5p |
| aga-miR-125-3p | aga-miR-79-3p | aga-miR-279-3p |
| aga-miR-1889-3p | aga-miR-9c-3p | aga-miR-996-3p |
| aga-mir-10368-5p | aga-miR-283-5p | aga-bantam-3p |
| aga-miR-10-5p | aga-miR-277-3p | aga-miR-1175-3p |
| aga-miR-iab-4-5p | aga-mir-10375b-3p | aga-miR-1891-5p |
| aga-miR-307-5p | aga-miR-965 | aga-let-7-5p |
| aga-miR-9b-5p | aga-miR-965-3p | aga-miR-12-5p |
| aga-miR-190-5p | aga-miR-124-3p | aga-miR-219-5p |
| aga-miR-276-5p | aga-miR-8-3p | aga-mir-932-3p |
| aga-miR-210-5p | aga-miR-993-5p | aga-mir-2c-3p |
| miR-iab-8-3p | aga-mir-998-5p | aga-miR-7-5p |
| aga-mir-10355-3p | aga-mir-285-5p | aga-miR-1-3p |
| aga-miR-1000-5p | aga-mir-31-3p | aga-miR-184-3p |
| aga-miR-305-5p | aga-mir-980-3p | aga-mir-31-5p |
| aga-miR-927-3p | chr2R_3959-miR* | aga-miR-72-1 |
| aga-miR-10-3p | aga-miR-276-3p | aga-miR-34-5p |
| aga-miR-1889-5p | chr2L_3426-miR | aga-miR-263a-5p |
| aga-miR-11-3p | aga-mir-2a-1-3p | aga-miR-281-3p |
| aga-miR-8-5p | aga-miR-2-1-3p | chrX_40848 |
| aga-miR-1-5p | aga-miR-13b-3p | aga-chrX_353716 |
| aga-miR-190-3p | aga-mir-2944a-1-3p | aga-miR-989-3p |
| aga-bantam-5p | aga-miR-92a-3p | aga-mir-999-5p |
| aga-miR-305-3p | aga-miR-137-3p | chrUNKN_30285 |
| aga-miR-iab-4-3p | aga-mir-932-5p | chr2R_12239-miR |
| aga-mir-980-5p | aga-miR-307-3p | aga-miR-981-3p |
| aga-miR-263a-3p | aga-miR-309-3p | aga-miR-133-3p |
| aga-miR-929-3p | aga-miR-3-1-miR | aga-miR-210-3p |
| aga-mir-252-3p | aga-miR-1174-3p | aga-miR-927-5p |
| aga-miR-263b-5p | aga-miR-275-3p | aga-miR-375-1-3p |
| aga-miR-315-3p | aga-miR-306-5p | aga-miR-315-5p |
| aga-miR-993-3p | aga-miR-14-3p |  |
| aga-mir-2944b-5p | aga-miR-970-3p |  |
| aga-mir-2944a-5p | aga-miR-125-5p |  |

**Table S2:**
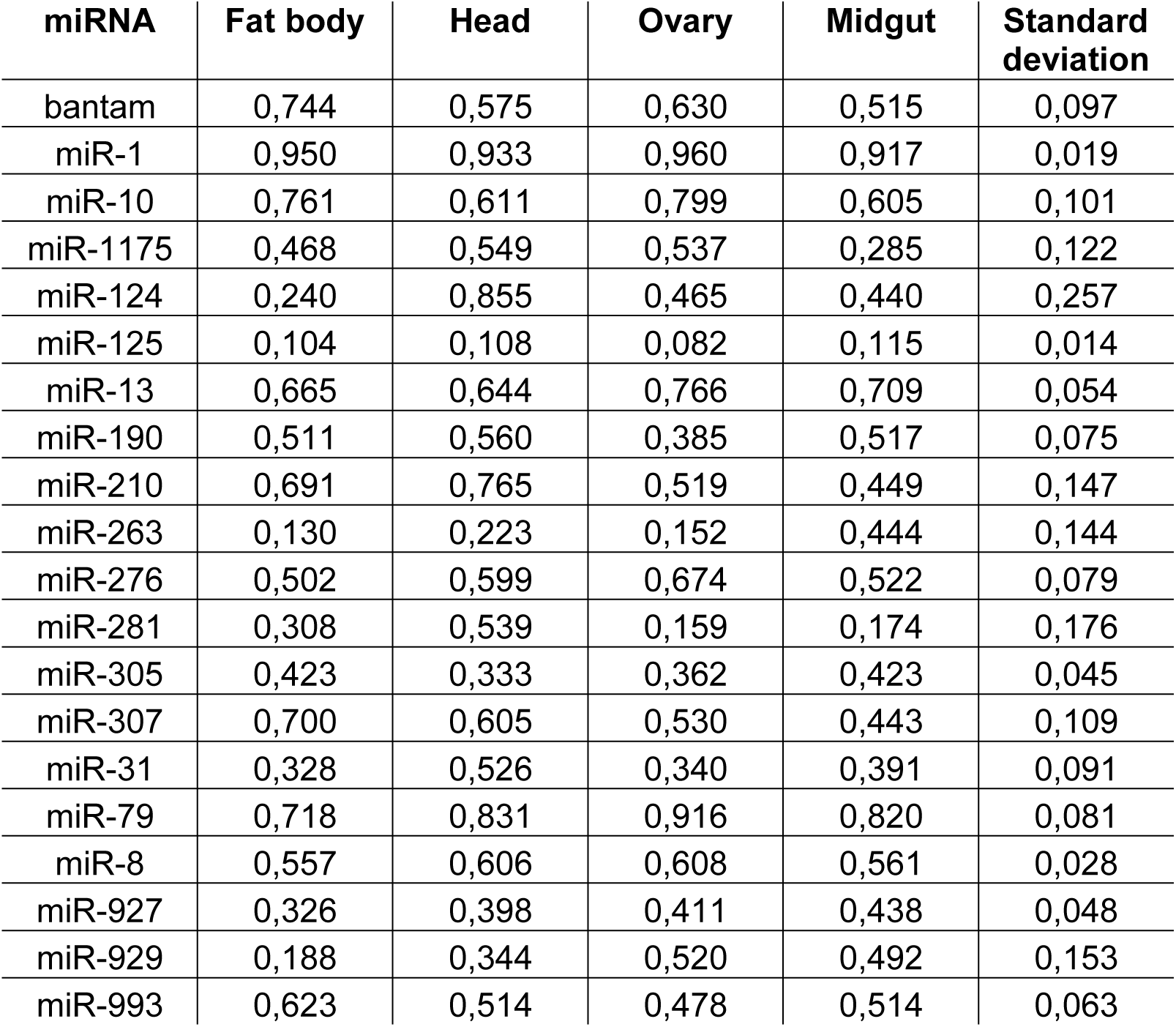
Proportion of miR-3p arm expression of miRNAs expressing both miRNA arms.

| miRNA | Fat body | Head | Ovary | Midgut | Standard deviation |
| --- | --- | --- | --- | --- | --- |
| bantam | 0,744 | 0,575 | 0,630 | 0,515 | 0,097 |
| miR-1 | 0,950 | 0,933 | 0,960 | 0,917 | 0,019 |
| miR-10 | 0,761 | 0,611 | 0,799 | 0,605 | 0,101 |
| miR-1175 | 0,468 | 0,549 | 0,537 | 0,285 | 0,122 |
| miR-124 | 0,240 | 0,855 | 0,465 | 0,440 | 0,257 |
| miR-125 | 0,104 | 0,108 | 0,082 | 0,115 | 0,014 |
| miR-13 | 0,665 | 0,644 | 0,766 | 0,709 | 0,054 |
| miR-190 | 0,511 | 0,560 | 0,385 | 0,517 | 0,075 |
| miR-210 | 0,691 | 0,765 | 0,519 | 0,449 | 0,147 |
| miR-263 | 0,130 | 0,223 | 0,152 | 0,444 | 0,144 |
| miR-276 | 0,502 | 0,599 | 0,674 | 0,522 | 0,079 |
| miR-281 | 0,308 | 0,539 | 0,159 | 0,174 | 0,176 |
| miR-305 | 0,423 | 0,333 | 0,362 | 0,423 | 0,045 |
| miR-307 | 0,700 | 0,605 | 0,530 | 0,443 | 0,109 |
| miR-31 | 0,328 | 0,526 | 0,340 | 0,391 | 0,091 |
| miR-79 | 0,718 | 0,831 | 0,916 | 0,820 | 0,081 |
| miR-8 | 0,557 | 0,606 | 0,608 | 0,561 | 0,028 |
| miR-927 | 0,326 | 0,398 | 0,411 | 0,438 | 0,048 |
| miR-929 | 0,188 | 0,344 | 0,520 | 0,492 | 0,153 |
| miR-993 | 0,623 | 0,514 | 0,478 | 0,514 | 0,063 |

**Table S3:**
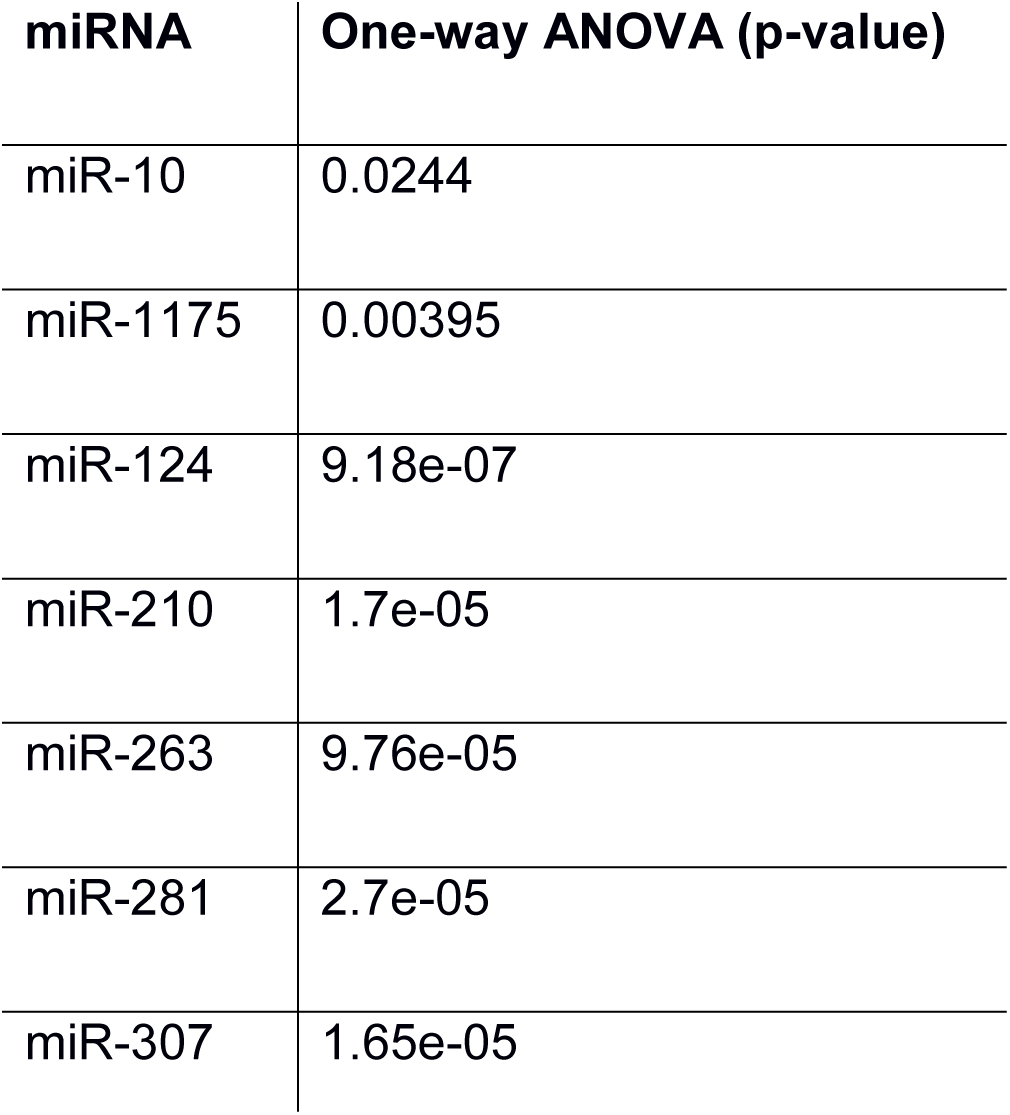
One-way ANOVA results for the differential expression of miRNA arm proportion across tissues.

| miRNA | One-way ANOVA (p-value) |
| --- | --- |
| miR-10 | 0.0244 |
| miR-1175 | 0.00395 |
| miR-124 | 9.18e-07 |
| miR-210 | 1.7e-05 |
| miR-263 | 9.76e-05 |
| miR-281 | 2.7e-05 |
| miR-307 | 1.65e-05 |

